# Real-time pooled optical screening with single-cell isolation capability

**DOI:** 10.1101/2023.09.21.558600

**Authors:** Praneeth Karempudi, Elias Amselem, Daniel Jones, Zahra Khaji, Maria Tenje, Johan Elf

## Abstract

In a pooled optical screen, a genetically diverse library of living cells is imaged and characterised for phenotypic variations without knowing the genotype of the cells. The genotypes are identified in situ after the cells have been fixed or by physical extraction of interesting phenotypes followed by sequencing. Mother-machine microfluidics devices are efficient tools in pooled optical screens since many strains can be imaged in the same field of view, but the throughput is often limited. In this work, we show a method to extract single bacterial cells from a compact 100,000-trap mother-machine-based fluidic device using an optical tweezer. Unlike previous devices, the fluids in our design are routed in 3D to enable fast loading of cells, increased trap density, and faster imaging. We have also developed software that allows real-time analysis of the phenotyping data.

## Introduction

Over the past two decades, microfluidic devices have been developed for numerous applications across biology, from DNA sequencing to rapid point-of-care diagnostics. Among quantitative cell biologists studying bacteria, so-called mother machine-based fluidic devices (Wang et al., 2010) are being used for various kinds of time-lapse microscopy. The fluidic chip usually contains a few thousand cell traps with a cross-section of 1x1 μm and a length of 20 to 50 μm. At the end of each trap, a bacterial cell is confined and allowed to grow and divide indefinitely with a fresh supply of growth media. These devices, combined with image processing software, provide an easy and powerful tool to observe cellular phenotypes over multiple generations.

Developments in our group (Baltekin et al., 2017a) have enabled fast loading and near-complete filling of all thousands of traps, making fast capture of bacteria and quick antibiotic susceptibility tests (AST) a reality. In the original closed-end mother-machine devices, centrifugation or a high concentration of cells (Thiermann et al., 2023; Wang et al., 2010) is needed to ensure proper loading. The fast loading capability of our chip is achieved by a constriction that stops the cells at the end of the trap but allows fresh media to pass over the cells. We use a pressure-driven flow controller to create a pressure difference between the fresh-media supply channel and the back channel. The pressure difference forces the cells into the growth traps, which makes loading of the chip fast and reliable.

Mother machines have played a critical role in studying phenomena related to replication initiation during the cell cycle (Knöppel et al., 2023; Le Treut et al., 2021), antibiotic resistance (Bakshi et al., 2021) (Brandis et al., 2023), mutation dynamics (Robert et al., 2018), and cell-to-cell variability in gene-expression (Kaiser et al., 2018), to name a few. However, studying phenotype-genotype relationships in large strain libraries using such devices is limited by our ability to recover the genotypes of the cells after phenotyping. Using fluorescence labelling strategies (Camsund et al., 2019; Kandavalli et al., 2022), we can determine the genotypes of all the library members in situ, but this strategy is impractical if only a few strains are of interest. Recently (Luro et al., 2019) published a device design with so-called Quake valves (Thorsen et al., 2002) added to the original mother-machine design (Wang et al., 2010). The valves make it possible to perform single-cell extraction from the device using an optical trap. The study takes an important step toward dynamic phenotyping of non-barcoded libraries, but still has some technical limitations. The published devices are limited to 16,000 cell traps spread over four independent units, each unit with its own connection ports to external fluidic reservoirs, which complicates operation. Moreover, the closed-end trap design makes sample loading more complicated. The cell segmentation and tracking are based on fluorescence images which limits the kinds of cell libraries that can be analyzed. In addition we have also added real-time image processing.

Advances in image analysis, especially deep-learning methods, have revolutionised bacterial cell biology. Software packages such as MM3 (Thiermann et al., 2023), Delta 2.0 (Cutler et al., 2022; O’Connor et al., 2022), and others (Jug et al., 2014; Sachs et al., 2016; Smith et al., 2019) are valuable tools when processing and analysing the wealth of data generated with the fluidic devices. Modern pipelines mostly depend on mixtures of neural-network models and classical algorithms for cell segmentation, cell tracking, and other fluorescence measurements, and use post-processing steps to curate for errors. Most of these software packages are built for post-analysis and are not designed with scalability, modularity, and fast acquisitions of large fields-of-views (FOV) as primary considerations. Imaging frequencies are adjusted to produce real-time-like effects (Lugagne et al., 2022).

Thanks to advances in genetic engineering and the powerful imaging and analysis software described above, studies of phenotype-genotype relationships in large-scale pooled libraries of cells (Camsund et al., 2019; Feldman et al., 2019; Lawson et al., 2017) are now often limited by the number of traps in the microfluidic chips and software upgrades to monitor real-time results. In most studies reported to date, the fluidic chip contains a few thousand traps, restricting the number of strains that can be observed with sufficient statistical power. When studying phenomena with high variance or rare events, obtaining good statistics gets even more challenging but the problem can be overcome by scaling device throughput.

However, scaling up the number of cell traps should be done without a corresponding increase in the number of external connections or inflation in the device footprint. Too many external connections will result in longer experimental setup time and an increased risk of contamination and bacterial growth at the connections. Contamination is particularly problematic if single-cell extraction is desired for further investigation. Increased physical dimensions of the device lead to extended travel times required for the microscope objectives, thus extending imaging time. To date, the largest number of traps reported for a mother-machine device is 16,680 (Luro et al., 2019). They use a closed-trap design which is relatively easy to parallelize, but the design does not allow medium to flow through the traps, which, e.g., makes loading slower. The open-end trap design facilitates loading but faces a topological challenge when scaled up. Each back channel introduced requires the addition of a new outlet port.

In this work, we present a new compact mother-machine design with 100,000 traps that is compatible with live-cell imaging and single-cell isolation. With this device, we expand the scope of single-cell phenotyping six times. To facilitate cell isolation, we have integrated pressure-controlled valves that separate the cell growth area from the clean cell isolation channels. To scale up the number of cell traps without increasing the number of outlet ports or device dimensions, we have introduced through-holes that allow fluid flow in 3D. These improvements are done without increasing the complexity of assembling the chips compared to most two-layer PDMS devices. The area of the chip used for imaging all traps is 1 cm x 1.5 cm, making it easy to use on modern microscopes with standard focus-locking mechanisms. We have also developed a real-time analysis pipeline with a graphical user interface for monitoring experimental progress and visualising the results.

## Results

### Fluidic design

We set out to develop a chip with 100,000 mother machine traps and only eight connectors while preserving attractive properties from previous designs, like fast loading (Baltekin et al., 2017b), and single-cell isolation capability (Luro et al., 2019). To achieve this goal, we had to route the fluids in 3D using a two-layer fluidic design (Thorsen et al., 2002) and through-hole fabrication in PDMS (Silva Santisteban et al., 2014). The two-layer design is shown in Figure 1a. The first layer has all the mother-machine cell traps and associated flow channels for cell loading, media supply, and waste. The second layer features the pressure-controlled valves and a separate flow channel system that is connected to the first layer via the through-holes (Fig 1b).

**Fig 1.**
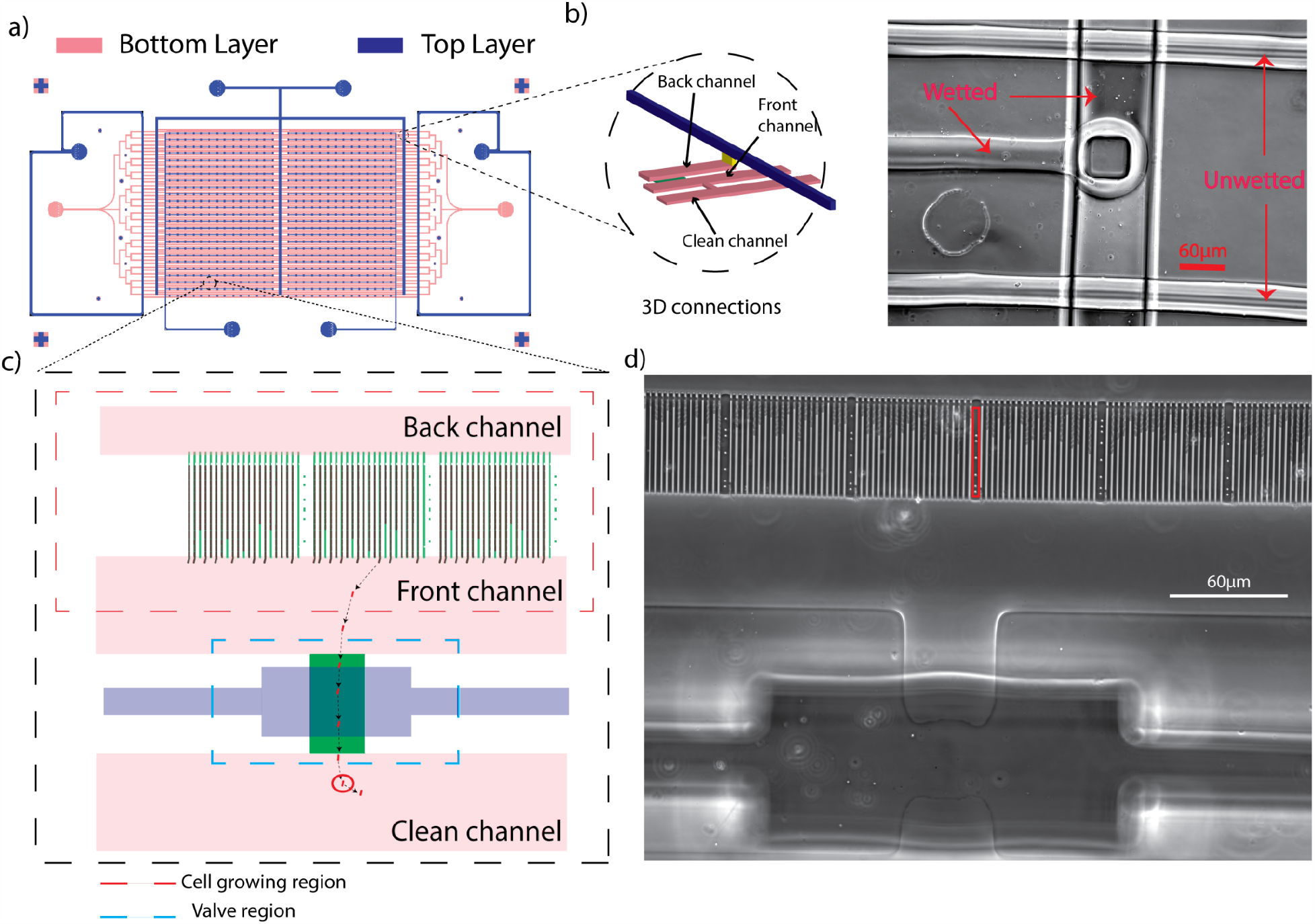
Fluidic design. a) Two-layer mother-machine chip with 100,000 channels. Bottom layer (pink) and top-layer (blue) of the chip. b) 3D connections present at the ends of bottom layer channels (left) and image of the 3D connections in 20x taken during the wetting phase of an experiment (right). The image shows one wetted channel and two non-wetted channels from the bottom layer (horizontal channels). A vertical channel runs in the second layer of the chip and is connected to a bottom channel in the middle with a through-hole made in PDMS. The image shows that fluids flow from top layer to bottom layer. The scale bar is 60 μm. c) A zoomed-in region of the chip showing cell growing region, valve region and clean channel (left). The front channel is connected to the clean channel with a connection in the bottom layer shown in dark green. The valve in the top layer,shown in blue, sits on these connections and is used for closing and opening the connections. d) A phase-contrast image of the chip imaged in 40x (right). The image shows *E*.*coli* cells loaded in the chip and a valve that is closed using a pressure of ∼ 1500 mbar. The red box in the image shows the unique barcode that separates sets of 21 mother-machine channels (a block). The scale bar is 60 μm.

The cell traps are arranged in 25 parallel rows, each row containing ∼4,000 mother machine traps, connected by front and back channels. Each row is split into two segments of 2,000 traps each. Each back channel has 3 through-hole connections, one at each end and one in the middle. The back channels of each row are connected to the second layer via these through-holes, providing independent back-pressure control. The mother-machine traps are grouped into sets of 21 traps (a block) with a unique barcode (Fig 1d) molded on the left and right of each set. This barcode uniquely encodes the position of each set on the chip and is used to keep track of trap numbers in the analysis software.

In-between the rows of cell traps run separate flow channels with periodic connections to the cell growth region at every 200 traps (Fig 1c). These connections can be opened or closed using the pressure valves in the second layer. These separate flow channels are connected to the second layer via 2 through-holes at the end of each row, providing independent pressure control. We call these channels clean-side channels as they are maintained separate from the cell-growing region during experiments (Fig 1c). To ensure the clean-side channels are completely isolated from the cell culture region, we follow the cell-loading protocol described in the methods section.

### Real-time software

To be able to identify cells with interesting phenotypes for isolation while an experiment is running, we have developed real-time analysis software. The real-time software also enables the identification of cells based on their dynamic phenotypes and closed-loop feedback control of experimental conditions. The cell segmentation, tracking, and growth-rate analysis pipeline runs on modest hardware resources in conjunction with image acquisition (Fig 2). Each task to be done in the pipeline is modelled as a sequence of operations handled by queues that are emptied after processing, preventing data accumulation. Unlike other image analysis pipelines (Thiermann et al., 2023), (Cutler et al., 2022; O’Connor et al., 2022), (Jug et al., 2014; Sachs et al., 2016; Smith et al., 2019), the analysis pipeline is bottlenecked only by microscope movement and focus stabilisation (∼500ms). All image processing operations are handled in less than 1s from acquisition, preventing the accumulation of data in memory. Data processing throughputs and number of traps processed are at least an order of magnitude more than those reported so far in literature for mother-machine devices. We also provide visualisation tools to monitor experimental results and make decisions on identifying potential traps to isolate cells from. For now, we only monitor the growth rates of single cells. Other computing operations, e.g. fluorescence measurements, can be added on top of the current analysis, without compromising performance or throughput as long as they run in ∼500ms per FOV acquired. The image acquisition sequence can also be modified based on the results of image processing, providing closed-loop feedback when required.

**Fig 2.**
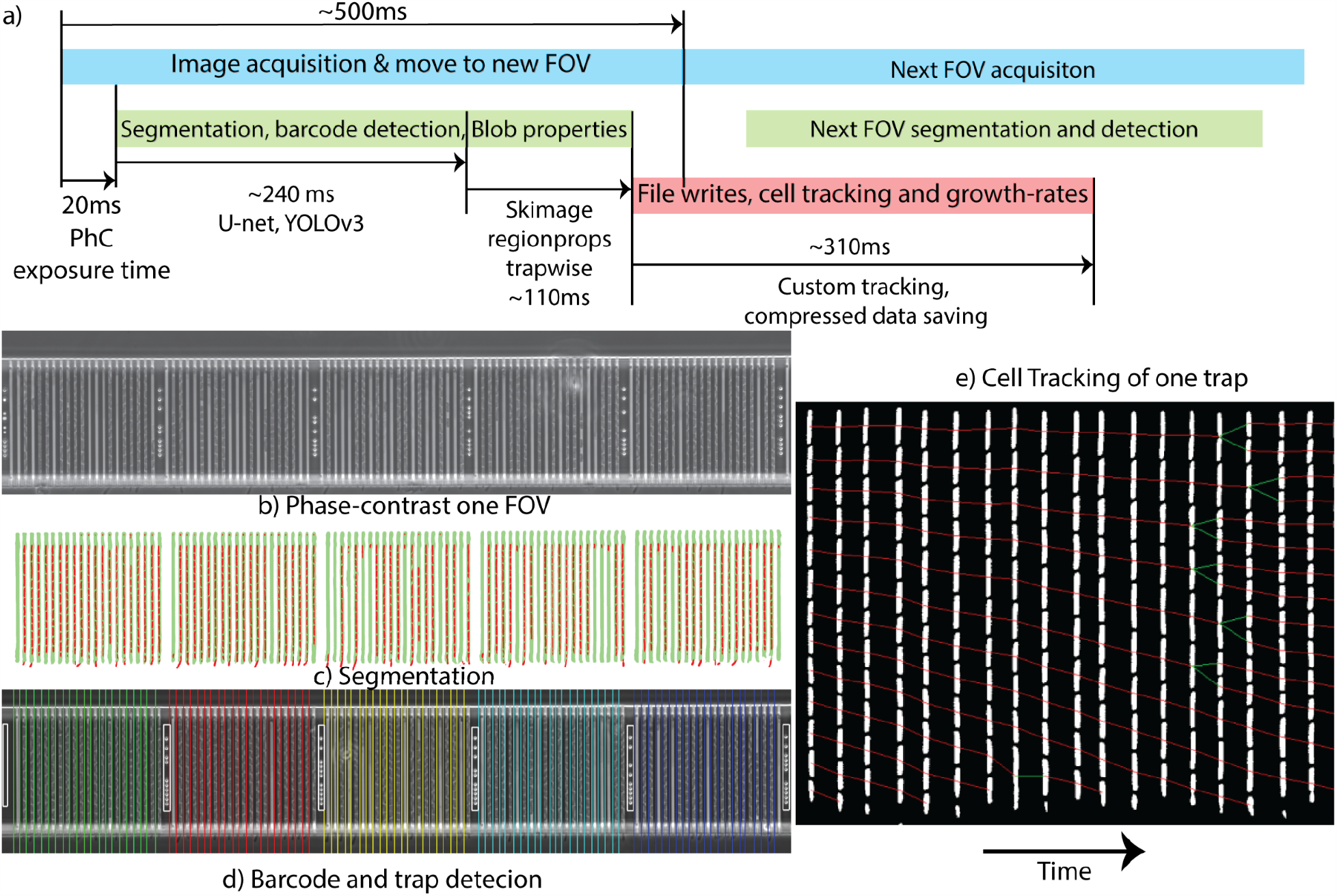
Real-time software. a) Image processing pipeline with timeline for analysing one FOV. The pipeline has three processes (blue, green, red) with queues between them. The first process (red) handles the microscope acquisition queue and passes images into the segmentation queue that feeds into the second process (green). The second process (green) analyses the phase-contrast images using neural networks (on GPU) and prepares data structures for cell tracking (on CPU) and storing. This data is passed onto the tracking queue. The third process gets data from the tracking queue and files from previous time points and does tracking of cells in one FOV trap-wise and writes results to disk. At the end of each process is a database and logging system step that manages data for visualisation in the GUI. b) Phase-contrast image of one FOV in 40x, 4096 x 900 pixels containing 105 traps. c) Cell and trap segmentation predicted by a single U-net overlayed. Cells are in red and traps are in green d) Detected barcode bounding boxes (white) and blocks of 21 traps identified, shown with lines of different colours next to the traps in that block. d) Tracking of cells in one-trap through time (5 min frame rate). Red links show cell-cell connection based on movement in the trap, and green links show connections that are divisions.

### Single-cell isolation using Optical tweezers

We use an optical trap to move cells with interesting properties from the front ends of the traps to the clean channel of the fluidic chip (Fig 3a-c). The cells are trapped at the focal point of a 50mW IR laser beam (1030 nm) and the microscope stage is moved to drag the cell. A spatial light modulator is used to create the laser focus at exactly the same imaging plane as the phase-contrast image. The valves connecting the cell traps with the clean area are opened only during the cell-isolation phase and remain closed during the phenotyping experiment. Once the cell is moved to the clean side, it can be collected into a tube or an agar plate for further investigation. A detailed protocol of the tweezing procedure is available in the methods section.

**Fig 3.**
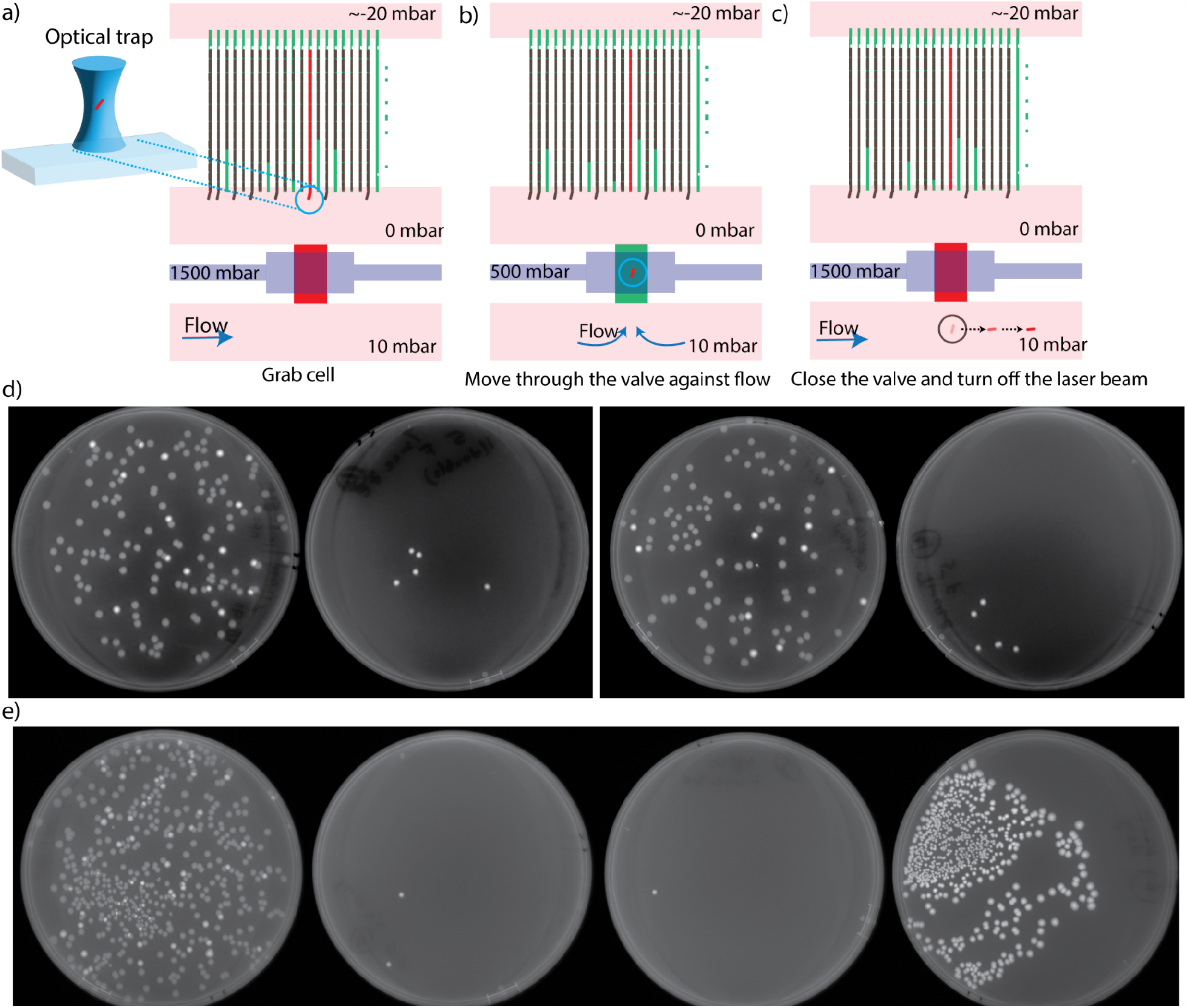
Single-cell isolation protocol and results. a) An optical trap (left) with 1030 nm laser used to grab cells at the end of trap. Relative pressures on the back channel (-20mbar), front channel (0 mbar) and clean channel (10 mbar) are maintained before grabbing the selected cell. The valve is closed at this time point using pressure (1500 mbar) in the top layer. b) After grabbing a cell, the valve is opened by reducing the pressure to 500 mbar allowing flow from the clean side to the cell-growing region. The cell is tweezed against the flow. c) The valve is closed once the cell reaches the clean side. At this point, the laser is cut off to release the cell. The cell runs to the collection port due to the flow. d) Collection plates from 2 control experiments. The plate on the left is a 1:1000 dilution of the cell culture mixture loaded into the chip. The right plate has colonies of fluorescence cells that were tweezed out of the chip. e) Collection plates for the third control experiment. From left to right, the first plate shows the 1: 1000 dilution of the cell culture loaded onto the chip, the second plate shows the collection plate after tweezing, the third plate shows the control plate after the tweezing step, the fourth plate shows the collection of cells that grew overnight at the outlet port.

To demonstrate single-cell isolation capability in this chip, we mixed fluorescent and wild-type *E*.*coli* 1:10 and repeatedly picked (n=5) fluorescent cells from different regions on the chip. The agar plates in Fig. 3D shows the distributions of cells in the loading culture (right) and the collection plate of the tweezing step (left). All control plates collected during the experiment to show non-contamination of the clean side of the chip are shown in SI section 1. The control experiment is repeated n=3 times and we show that we can reliably flush the selected cells out of the chip. Out of the 15 cells picked in 3 experiments, 13 were collected on the plates and 2 were likely stuck in the collection port. Fig. 3e shows a plate with cells collected from overnight growth at the collection port being all fluorescent. All cells extracted from the chip with the optical tweezer were fluorescent. The time from the point of picking the cell at the end of the trap to releasing it on the clean was ∼40s. Video recordings of the picking process are available as Supplementary movies.

## Discussion

We have made two innovations that will facilitate large optical pooled screens in the future. First, by routing the back channels in 3D we can keep the number of fluidic connections and the physical footprint of the device manageable, despite scaling to 100,000 cell traps. At the same time, we keep the functionality with a clean channel where cells are moved with the optical tweezer to be extracted from the chip. Secondly, we have developed a real-time cell segmentation and tracking algorithm that makes it possible to stop acquisition and start tweezing cells when sufficient data has been collected.

The capability to isolate one strain out of 100,000 different strains based on dynamic phenotypic traits opens for a plethora of possible screens. A few that come to mind are altering promoter or operator sequences and screening for dynamic responses in gene expression following a chemical alteration, or changing the protein-coding sequences and screening for functional readout of the protein such as regulatory, enzymatic, and structural properties. It is also possible to make many different genetic perturbations using CRISPRi or transposons and screen for dynamic changes in biological processes.

## Methods

### Bacterial strains and cell-culture conditions

Control experiments use two strains of *E*.*coli*, of which one expresses fluorescence (Eco MG1655 rph+ DELfliA::kanR intC::P70-venusFast CmR) whereas the other one does not (Eco MG1655 rph+ DELfliA::kanR). Cells are grown in LB + 1:2000 kanamycin (50 mg/ml stock) overnight at 30 °C and diluted next day in M9 succinate + amino acids (RPMI 1640 diluted 1x, R7131; Sigma-Aldrich;) + Pluronics (F-108; 542342; Sigma-Aldrich; 0.085% (wt/vol) final concentration). After 4 hr at 30 °C, different proportions of cells are mixed before loading them on the chip. Training data for segmentation algorithms was obtained using *E. coli* (K12 MG1655 intC::P70-venusFast) imaged in fluorescence and phase contrast at both 100x and 40x.

### Optical setup

For imaging and tweezing we use a Nikon-T2 E microscope with a perfect focus module (PFS-MP) which allows focus lock while tweezing. The microscope is equipped with a 20x phase contrast objective which gives a good overview of the traps. 20x phase contrast is used to check that all valves are fully closed during the wetting of the chip. A high numerical aperture (NA of 1.3) 40x phase contrast objective (Nikon MRH01401) is used for single-cell imaging and tweezing. For imaging fluorescence, an epi configuration is used in combination with the light source (Lumencor Spectra III Light Engine), filter set (dichroic: Semrock Di02/R514, excitation; Chroma Z514/10, emission; Semrock FF01-542/27), and camera (Imaging source; DMK 38UX304). The tweezer is a custom-made system with a 1W fiber laser (MPBC 2RU-YFL-P-1-1030) operating at the wavelength of 1030nm. The beam is expanded onto a spatial light modulator (Hamamatsu X15213-03). By producing a blazed grating pattern (grating with a sawtooth profile) (Ritsch-Marte, 2017) on the Spatial Light Modulator (SLM), the first-order refraction is isolated and routed to the back focal plane of the objective using a dichroic mirror (Thorlabs DMSP900R). To protect the camera from reflections and stray tweezing laser light, a bandpass filter (Thorlabs FESH0750) is inserted in the camera path. Using the SLM, we can fine-tune the focal spot to be at the same plane as the imaging plane which is where the tweezing is taking place. The tweezing power used is about 50mW measured after the objective without mounting a sample. All images are acquired with Micro-manager 2.0 directly or by using Pycromanger v0.20.0 to interact with the microscope through a custom pipeline built in Python. Each FOV imaged in phase contrast results in an image of size ∼4096 x 1024 pixels, where each pixel corresponds to 86.25 nm at the imaged resolution.

### Wafer fabrication

Two silicon wafers were used to fabricate the PDMS chips. SI Fig 3 shows the outlined fabrication sequence for both wafers. The first wafer contains structures fabricated at 3 different heights. First, a SiO2 layer of height 1250 nm is grown on a 4-inch Si wafer (University wafer, P<100> 525 ± 25μm) in an oxidation furnace (Koyo Lindberg, Micro TF-6) for 7.5 hrs using wet oxidation process. This layer is etched later to form structures for the 100,000 cell traps.

Positive PMMA resist (PMMA4, MicroChem) was spin-coated to 100 nm thickness and cell traps were patterned using e-beam lithography (NanoBeam Ltd, nB5). PMMA was developed in 1:4 MIBK (Microchemicals) to Isopropyl alcohol for 2 mins and rinsed in Isopropyl alcohol for 45 s. After e-beam lithography, 30 nm of Cr was deposited on the wafer by e-beam evaporation (Kurt J. Lesker Company, PVD 75) and unexposed PMMA was washed away in Acetone. Using the Cr layer as a hard mask, the SiO2 layer was etched using dry etching (CF4 - 40 sccm flow, 100 mbar pressure, 100 W power) in a reactive-ion etching chamber (Oxford Plasma Technology, Plasmalab 80 Plus) to leave only the cell-trap structures. The wafer was cleaned in Piranha solution (2 : 1 H_2_SO_4_ : H_2_O_2_ solution) for 15 mins, to increase the adhesion of the following SU-8 resist layers. Negative SU-8 3025 resist was spin-coated onto the wafer to form a 50 μm thick layer. This layer was used to pattern the through-holes connecting the two fluidic layers and the resist was patterned into pillars located at the ends of back channels and clean channels using UV lithography (365 nm, MA6 Mask Aligner, Karl Süss). The wafer was baked post UV exposure (20 seconds at 12 mJ/cm^2^) on a hot plate at 65°C for 1 minute and 95°C for 5 minutes. The wafer was then developed in mr-Dev 600 (Micro resist technology GmbH) to remove non exposed resist. The wafer was then hard baked on a hot plate with a slow temperature-ramping from 95°C to 175°C over 40 min to increase the cross-linking of the SU8 pillars. Next the wafer was vapour primed with Hexamethyldisilazane (HMDS) in a Star 2000 oven to increase the adhesion of the last layer of photoresist. Positive resist (AZ 10XT 520cP, Microchemicals) was spin-coated to a thickness of ∼10 microns. The AZ 10XT resist was exposed to UV light to define the flow channels that supply media to the cell traps and the non-exposed resist was developed (AZ 400K 1:4, Microchemicals) before the wafer was carefully rinsed in water. Resist reflow was used to curve the 10-micron channels at 130°C for 10 minutes on a hot plate to aid complete closing of the valves when the wafer was moulded into PDMS. Finally, a thin coating of C_4_F_8_ was deposited using deep reactive ion etching (DRIE) (PlasmaTherm SLR) to make the surface superhydrophobic.

For the second wafer, all structures were made in a single layer of SU8-100 resist spin-coated onto a Si wafer to reach ∼100 μm height. The second wafer design was adjusted and scaled with an additional 1.25 % in all dimensions to account for PDMS shrinkages in the replica-molding step of chip construction, described below.

SI Fig 3 shows the fabrication steps and design patterns structured in each step.

### PDMS chip molding

Two-layer chips were made by bonding two partially cured PDMS layers (SYLGARD™ 184). 1:20 proportion of curing agent to base was used for spin-coating PDMS on the first wafer at 3,700 rpm for 30 s followed by 4,200 rpm for 30 s, resulting in an even layer with an approximate height of 40 μm. After spin coating, the wafer was allowed to sit on a flat surface for 10 min for the PDMS to settle down on the structures before the PDMS was cured at 80°C for 15 min in an oven. 1:5 proportion was used for molding structures on the second wafer. After depositing PDMS onto the second wafer, it was degassed in a chamber under vacuum to remove air bubbles, before it was cured at 80°C for 22 minutes. Different curing times were used for the two layers to reach appropriate levels of stickiness, by adjusting previously identified curing times. Chips were cut from the second wafer’s PDMS cast and placed on the first wafer to manually align the pressure values and the 3D through-holes with the structures on the bottom layer of the fluidic design. At this stage, it is important to apply gentle pressure on the chip to get rid of any air pockets trapped between the two layers and ensure close contact between the two PDMS layers. After alignment, the two layers were fully bonded in an oven at 80°C overnight (16 hours). Finally, the two-layer chip was cut out into an appropriate size. Inlet ports were punched and the chip was bonded onto a clean glass coverslip using air plasma (Henniker Plasma HPT-100, 50% power, 1 mbar pressure, 30s). The bonded chip was incubated in an oven at 80°C for one hour. It is critical to avoid dust at all stages of chip molding, especially after spin-coating, during layer alignment, and bonding to the coverslips.

### Cell loading and flow preparations

The fluidic chip is connected to external fluid reservoirs using Tygon tubing (i.d = 510 μm, o.d = 1,524 μm; Saint-Gobain), custom-made metal pins (23-gauge, 14-mm-long tubing bent in the middle at 90°; New England Small Tubing) and 15 ml Falcon tubes with reservoir adapters (Elveflow, Size S), where the flow is generated by a pressure-driven flow controller (Elveflow O1Mk3+) to pressurize fluids in 15 ml falcon tube reservoirs.

All tubes and adapters are cleaned in ethanol and water twice before use to ensure no contamination. The metal pins used to interface with the chip are autoclaved and sonicated in 96% EtOH for 6 hrs and dried before use.

In total, we have 6 falcon tubes connected to the chip during the wetting phase of the experiment: one tube for the valves, one for the back channels, 2 for the front channels, and 2 for the clean-side channels (SI Fig 2). When we refer to pressures on a particular channel in the chip, we refer to the pressures applied on the fluid reservoirs connected to that channel.

Before we wet the chip, we pressurize the valves with culture media slowly from 0 to 1500 mbar and they are inspected at 20x to ensure all of them are closed. Usually, at around 1000 mbar most of the values are closed. Then we wet both the clean side ports with culture media at 200 mbar. If the valves closed completely, there should be no way for fluid to reach the cell-growing side. We can notice any potential leaks to the growing side with the human eye at this stage of the experiment.

Back channels, media, and cell-loading ports are pressurized with culture media until the chip is completely wet. One of the clean side port connections is swapped for a smaller tubing of length < 10 cm and is used for collecting single cells on Luria Agar plates. Once the chip is completely wet, cells are loaded from the cell-loading port at 300 mbar for about 10 min until most traps are loaded, after which it is dropped to 0 mbar while increasing the pressure on the media port to 400 mbar simultaneously. Both back channels and cell-loading tubes are lowered in height and non-pressurized as they are no longer necessary after loading cells.

### Tweezing protocol

To ensure a clean collection of cells, it is critical to control the pressures on all ports carefully as the tweezing force is weak compared to normal flow forces in the chip. To prepare for tweezing cells, the four tubes connected to the chip are arranged in the following decreasing order of height: clean side left tube, fresh-media tube, back-channels tube, and cell-loading tube. This ensures that cells growing out of the traps are collected in the cell-loading tube, while maintaining enough pressure difference between front channels and back channels to keep cells in the traps.

To start tweezing, the IR laser is turned on and the spot at which the beam hits the focal plane is marked with a circle in micromanager ImageJ, before pushing a long-pass filter (750 nm) into the camera path. This helps us visualize the cell trapped in the center of the circle, without seeing the laser beam spot. An epi-shutter is used to cut off the laser when required and is used to control when the beam is active on the cell being trapped.

Once we decide which trap to pick the cells from, pressure on the media is lowered from 400 mbar to 25 mbar, and pressure on the clean side is increased to 10 mbar. This ensures that cells stay in the trap and all fluids are directed towards the back channels and the cell-loading port. When the valves are opened there is enough flow to keep any cells from crossing the valves other than the one we are tweezing. Pressure on the valves is reduced to 500 mbar to open them partially while tweezing. The valves are closed after the cell has been transported to the clean side. The cell is then released from the optical trap by closing the epi-fluorescence shutter. It is important to hold onto the cells in the optical trap until the valves are closed to guarantee that collected cells don’t escape to the cell culture side due to the flow. A typical process from trapping a cell on the growing side to releasing it on the clean side takes 40 s.

### Cell segmentation

Phase-contrast images obtained with the 40x objective and the camera contain ∼ 105 cell traps each and has a size of typically 4096 x 1024 pixels. These images are segmented using smaller variants of U-nets (Kandavalli et al., 2022) trained on 40x and 100x images using methods described before (Kandavalli et al., 2022). The smaller U-net has fewer convolution-layer channels and is trained to predict both cell and trap segmentation masks in one inference step unlike models described before. The inference time on one image of size 4096 x 1024 is ∼250 ms using a computer equipped with NVIDIA RTX3090 24GB GPU, Intel i9 processor and 32GB DDR4 RAM). Data annotation, loss functions, and training steps are similar to those described in (Kandavalli et al., 2022)

### Barcode detection and Data storage

Barcodes are detected using a YOLOv3 model (Redmon and Farhadi, 2018), trained on 1200 annotated images. Bounding box predictions are corrected for errors using highest probability prediction boxes. The barcode model runs in ∼15 ms on 4096 x 1024 pixel images. Combining both trap segmentation and barcodes, we aligned traps from two timepoints. Segmentation masks, trap locations, and barcode locations data are then written to disk with compression and data-access patterns optimized for fast reads and writes using Zarr-Python.

### Cell tracking and growth rate calculations

Cells between frames at two timepoints are linked based on overlapping scores using a priority strategy similar to those described before (Ruzaeva et al., 2022). The priority for linking cells is based on one-dimensional overlap scores between segmented cell masks. The links between cells are corrected during assignment for rapid changes in area and center-of-mass movement, to account for errors in previous segmentation steps or noisy overlap scores. Growth rates are also calculated during linking cells as a percentage change of area between the connected cells. All trajectories are written in JSON format (JSON, 2005) to be used later for visualization in the graphical user interface (GUI).

### Running real-time experiments

Each field-of-view (FOV) of 40x phase-contrast imaging contains ∼ 105 traps and a total of 1000 FOVs are required to image all 100,000 traps. XYZ stage locations of FOVs are calculated automatically using interpolation between corners of the rectangular region containing the traps. All image processing and microscope hardware control is handled by real-time software tools.

Real-time software comes with 4 tools. One for running experiments (runner), one for viewing results (viewer), one for debugging results on live-mode, and a final one for defining FOVs to image on the larger chip area. All 4 tools are controlled using a single .yaml containing all the parameters required for setting up experiments. The image acquisition sequence can be programmed by specifying micromanager configuration presets and exposure times for acquiring required images. The positions on the chip are arrayed based on different movement patterns and corners marked on the imaging area. Variations in z (focus) are interpolated between the corners using bilinear interpolation and are adjusted manually before starting acquisitions. The viewing system can be used to monitor the progress of different queues in the processing pipeline to ensure no data accumulation in memory. It can also be used for monitoring data such as growth rates and lineages from any specific cell-trap of the data acquired. Identification of interesting phenotypes can also be done in the viewer. This will help to filter for positions and cell-trap locations from which we can tweeze cells. Overall the real-time system consumes ∼11 GB of GPU RAM, ∼10 GB of CPU RAM and ∼ 30% of available CPU cores (6 out of 20). All code is written in Python3 and is packaged to be installed using Python-Pip3 distribution infrastructure and custom conda-environment configurations with appropriate dependencies.

## Acknowledgements

We wish to thank I. Barkefors for helpful input on the manuscript. We acknowledge funding from SSF (ARC19-0016), the European Research Council (BIGGER:885360), the Knut and Alice Wallenberg Foundation (2016.0077; 2017.0291; 2019.0439) and the eSSENCE e-science initiative to J.E.as well as SciLifeLab postdoc grant to A.K. We acknowledge Myfab Uppsala and Myfab KTH for providing facilities and experimental support, especially Örjan Vallin for the support with e-beam lithography. Myfab is funded by the Swedish Research Council (2019-00207) as a national research infrastructure. We also thank Uppsala Antibiotic Center (UAC) for funding part of the project.

## Author contributions

J.E conceived the project. P.K designed and fabricated Si-molds in the clean room under supervision of M.T. P.K. wrote all the code for the real-time system and performed all the control experiments. E.A built the optical tweezer setup.

D.J. performed cloning and strain construction.

## Code and data availability

All code will be available at https://www.github.com/karempudi/sting upon publication. All raw data and fluidic chip designs will also be made available upon publication.

## Supplementary Information

### Control experiments

During the control experiments, media is collected from the clean side exit port of the chip at various time points. First ∼100 uL of media is collected on two different plates after wetting the chip and after loading cells (SI Fig 1a, 1b). No colonies on these plates indicates that the clean-side has no contaminants before the start of the tweezing step.

When the valves are opened in the chip, all valves on all the rows of the chip are opened at the same time. To check for possibilities of cells crossing the valves when they are opened, we simulate the valve opening and closing process for a tweezing experiment. The pressures on the front channel, back channels and the clean channel are adjusted to values used for tweezing and the valves are opened for 1 min and then closed. Media is flushed in the clean channel for 2 min at ∼200 mbar to collect any contaminants that might have crossed the valves. This process is repeated 5 times. No colonies on this simulation plate indicates that no cells crossed the valves without tweezing (SI Fig 1c).

5 cells expressing fluorescence were identified on the chip after imaging in the YFP channel and were moved to the clean side using the protocol described in (Main Fig 3a). Media was flushed for 2 min at ∼ 200 mbar after every move of the fluorescent cell to the clean side. After tweezing cells, media is flushed in the clean-channel for 15 min at ∼200 mbar to collect all the cells on the plate (SI Fig 1d). After tweezing cells, we also collect an extra plate by flushing the clean channel at ∼200 mbar for 15 mins (SI Fig 1e). This is to ensure that any cells that weren’t collected in the tweezing plate would be collected in this plate.

**SI figure 1.**
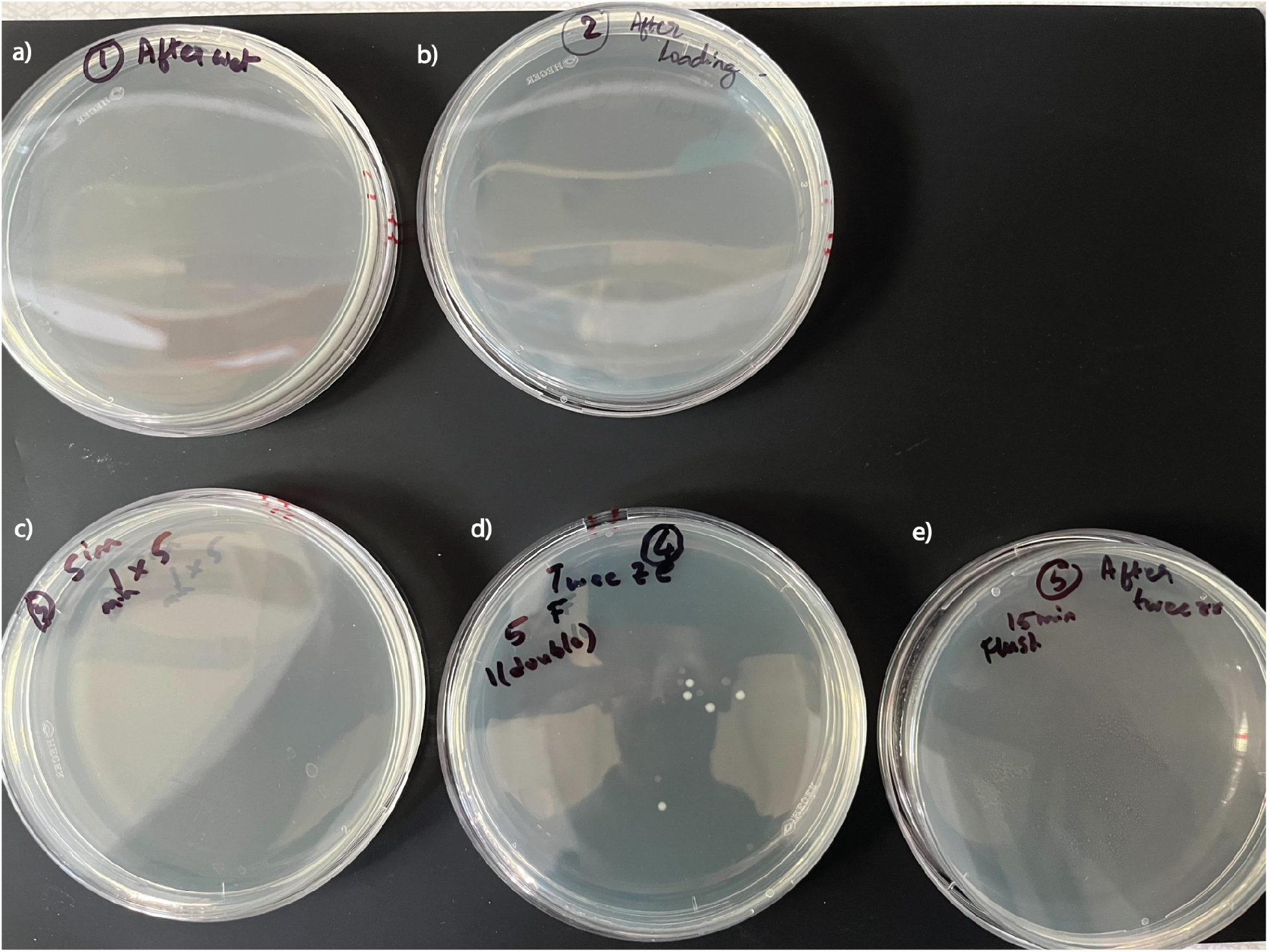
All plates collected during control experiments. Each plate has ∼100 μl of media collected from the clean channel outlet port at different time points during one control experiment a) After wetting the chip b) After loading the cells c) Simulation of opening of the valves for 1 min 5 times, with fluid collection for 2 min at 200 mbar after for every opening. d) 5 cells tweezed based on fluorescence observation collected on the plate. Media was flushed for 2 min at 200 mbar after tweezing a cell and for 10 mins at 200 mbar after all 5 cells were tweezed. e) After tweezing we flush the clean side for 15 mins at 200 mbar to check for any cells that didn’t come out onto the tweezing plate in SI Fig 1e.

**SI figure 2.**
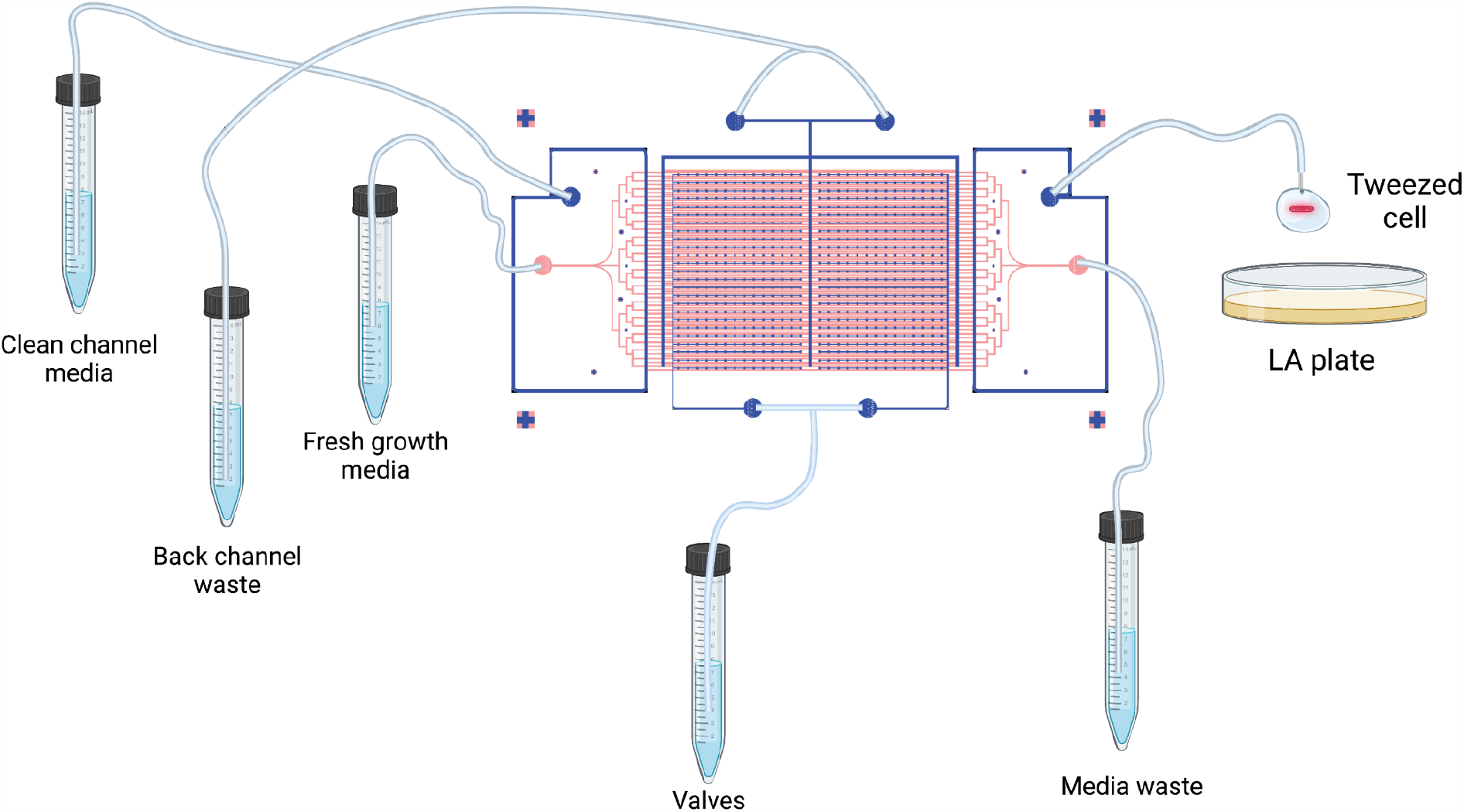
Fluidic connections and ports. A wiring diagram showing fluidic connections, tubing and collection of a cell on an agar plate. The figure is not drawn to scale, for illustration purposes only.

**SI figure 3.**
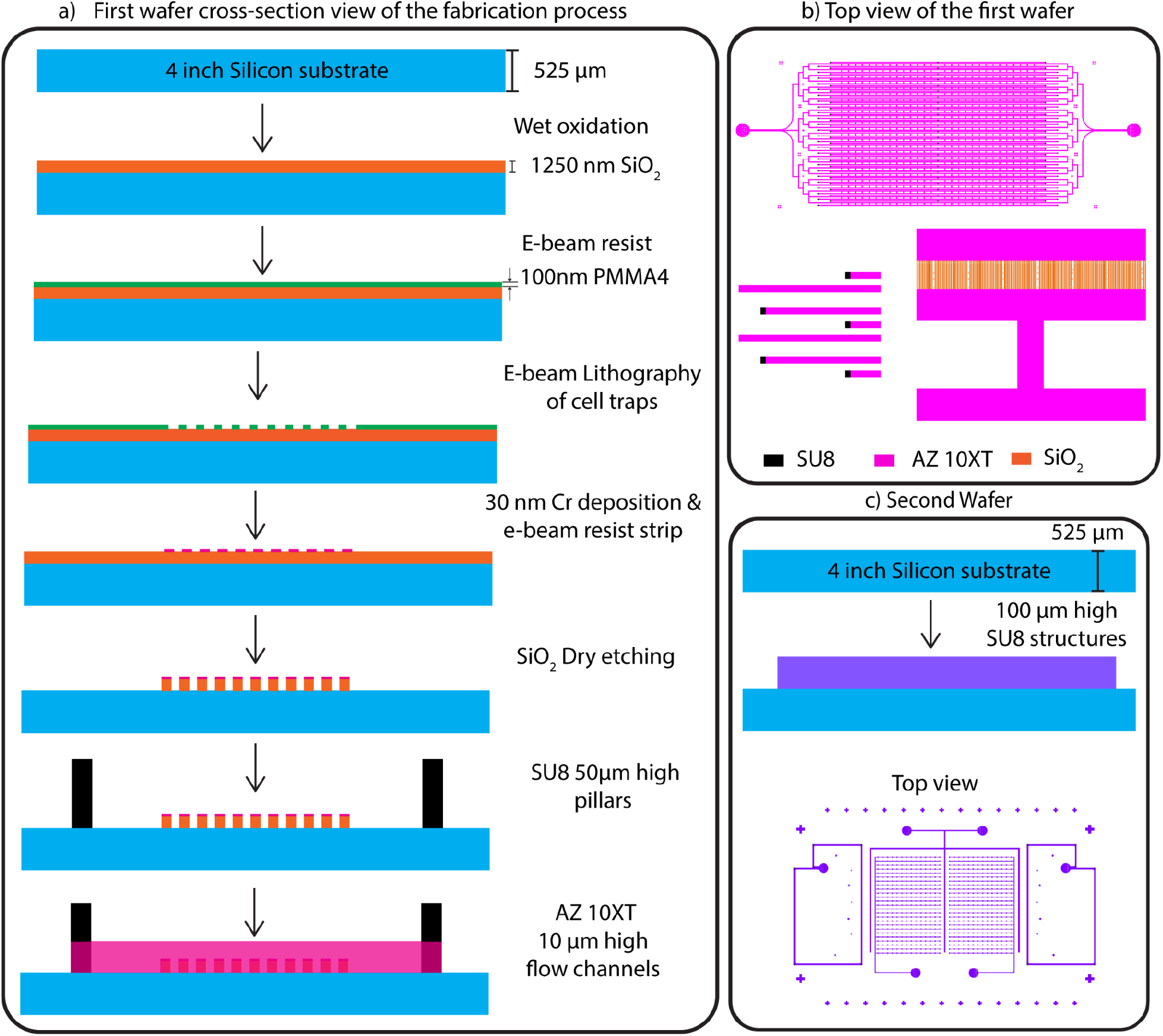
Silicon wafers fabrication steps. a) Fabrication steps showing the cross-section of the first wafer. Silicon substrate is shown in blue, SiO_2_ layer in orange, SU8 layer in black and AZ 10XT in pink. Features are drawn for illustration purposes and are not drawn to scale. b) Top view of the features on the first wafer. Mother-machine traps are in orange (1250 nm high SiO_2_ layer), flow channels (10 μm high AZ 10XT) and pillars (50 μm high SU8 layer) c)Fabrication steps for the second wafer showing a Silicon substrate and a single SU8 layer of 100 μm thickness. The top view shows the pattern transferred to this layer using UV lithography.

